# Status of urban feral cats *Felis catus* in England: A comparative study

**DOI:** 10.1101/394643

**Authors:** Nicholas P. Askew, Flavie Vial, Graham C. Smith

## Abstract

This study sought to determine whether a change in the abundance of feral cats *(Felis catus*) in three areas of England had occurred between the completion of a survey undertaken by the Ministry for Agriculture Fisheries and Food in 1986/7 and the turn of the century. In the event of a rabies outbreak occurring in Britain, feral cats would be one vector of the disease that would need to be controlled under the Rabies (control) Order 1974. A total of 741 “high risk sites”, found to provide appropriate conditions for feral cats, were surveyed between 1999 and 2000. The total number of feral cat colonies located within the survey areas was found to have fallen by 37% from 68 in 1986 to 43 in 1999/2000, translating to an estimated 212-247 fewer individual feral cats. Factories/trading estates and industrial premises continued to be the most common sites associated with urban feral cat colonies. However, the closing down of many traditional industries, such as mills and dockyards, and their replacement by more secure and insulated modern buildings, less amenable to feral cats finding warmth and food, had assisted the observed fall in numbers along with the effectiveness of neutering programs which are now taking place on many sites. Through this study information regarding feral cat colonies’ in urban landscapes as well colony size was gathered and fed into rabies contingency plans to help keep Britain rabies free into the future.

## 1. Introduction

At present, domestic cats *Felis catus* are the most abundant carnivore in Great Britain and their numbers appear to be growing (Woods et al., 2003). The national domestic population is estimated to have increased from 6.0 million cats in 1982 (Rees, 1982), to around 10 million in 2006 (Murray et al., 2015). Despite a long period since first being domesticated in Egypt around 1600 BC (Beadle, 1977; Todd, 1978), the cat has retained its natural ability to be a self-sufficient predator. Consequently, domestic cats readily revert to a free-living, feral status (Tabor, 1981) and can form colonies which hunt and scavenge for their food (Dards, 1978b; Macdonald and Apps, 1978; Dards, 1981). Harris *et al* (1995) estimated the number of feral or semi-wild cats in rural areas of Great Britain to be 813,000, but several studies have noted an unknown numbers of feral cats were also found in loose association with hospitals, industrial premises, residential areas (Rees, 1981; Harris et al., 1995) and dockyards (Dards, 1978a; Page et al., 1992) in urban areas.

Globally, feral cats have been well studied, with the majority of work focussing upon their behaviour and social interactions (Dards, 1978b; Macdonald and Apps, 1978; Apps, 1981; Dards, 1981) and methods for controlling them as a pest species (Hammond, 1981; Kristensen, 1981; Rees, 1981; Remfry, 1981). However, little data is available regarding the abundance of feral cat colonies within urban areas despite their potential for constituting a significant threat to public health, maintaining and transmitting the pancreatic fluke (Carney et al., 1970), leptospirosis (Ferris and Andrews, 1965), toxoplasmosis (Dubey, 1973) and acting as a vector of rabies between wild animal populations and humans (Bunn, 1991). As a result of these concerns the presence of feral cat colonies has been actively discouraged on some sites, with control being carried out on many groups. The most effective method of control developed was to trap, neuter and release the cats, limiting their ability to reproduce and resulting in a steady decline in the number of cats (Hammond, 1981; Kristensen, 1981; Rees, 1981; Remfry, 1981).

The United Kingdom has essentially remained free from rabies since 1903 following the implementation of strict anti-rabies control measures, including: stringent import controls, compulsory quarantine requirements, vaccination of imported cats and dogs and server penalties for offenders (Fooks et al., 2004). Even with these tough measures in place, sporadic cases of imported animals infected with the virus have been reported (Fooks et al., 2004). However on parts of mainland Europe, rabies is still prevalent, with 1014 wildlife (excluding bats) cases and 1900 domestic animal cases still being reported in 2017 (WHO, 2018). In the event of an infected animal coming into contact with another unconfined individual, the Rabies (control) Order 1974 provides wide ranging powers for dealing with this, including the declaration of an infected area and detailing control measures that may take place within them (Nicholson, 1981). Within such an infected area, it would be the Department for Environment, Food & Rural Affairs’ (DEFRA) responsibility to control any straying or free-living animals to prevent the spread of the virus.

The definition of a feral cat colony used for rabies contingency purposes is:

*“A group of three or more un-owned, un-wanted and un-confined cats, Felis catus, which the owner of the property where they occur would be unable to confine, if required to do so under the Rabies (control) Order 1974”*(Page and Bennett, 1994).

A prerequisite for control is locating the sites most frequently associated with such cat colonies and having accurate data available on expected densities within an urban area. For this reason between 1986 and 1987 three surveys were undertaken in separate areas of England by staff from the Ministry for Agriculture Fisheries and Food (MAFF- predecessor of DEFRA), to determine the locality and abundance of feral cat colonies (Page, 1986c, b, a). Here, we present the results of an identical survey undertaken between 1999 and 2000 and compare our results to the previous work. We aimed to assess the status of feral cat colonies found within the three areas, and to highlight any changes in their locations or densities that may have taken place to inform rabies contingency planning.

## 2 Methods and Study area

The surveys were undertaken in three industrial areas of England, Oldham, Swindon and the Wirral (Figure 1), originally selected by MAFF because they were thought to contain many of the urban habitats occupied by feral cats within the United Kingdom as described in Rees (1981).

**Figure 1:**
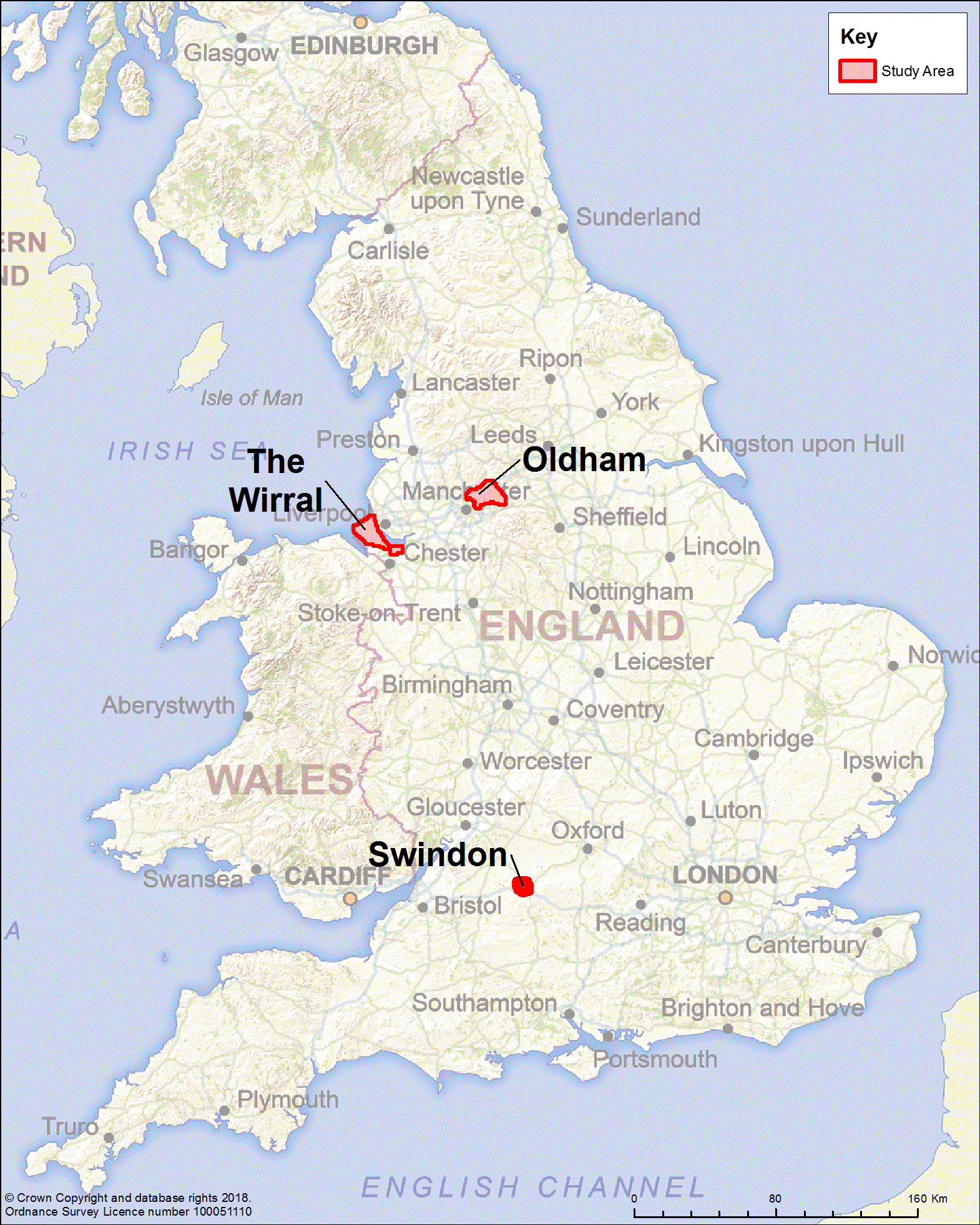
Map of England showing location of study sites.

The metropolitan borough of Oldham is situated within Greater Manchester in northern England. The borough was built up around the cotton-spinning and weaving industry, although the number of mills has today declined and many of the old large buildings stood derelict during the surveys. The study area was 136 km^2^ and encompassed Oldham and Saddleworth Moor and was bounded to the north by the M62 motorway.

Swindon is situated in Wiltshire, southern England, and was historically the home of the Great Western Railway Works before its closure in 1986. Since them, the town has undergone extensive redevelopment with the building of several large industrial estates. The study area was 41.5 km^2^ and encompassed the urban limits of the town, being bisected east to west by the railway line.

The Wirral peninsula, Merseyside, is situated in north-west England. It projects out into Liverpool bay, is bounded to the west by the estuary of the river Dee and to the east by the river Mersey. Ellesmere Port is found at the southern end of the study area and contains numerous large chemical plants, manufacturing premises and extensive petro-chemical works. The survey area was 228 km^2^ was largely built-up and home to a large ship-building industry.

The previous study boundaries were located based on simplified maps found in the report for Oldham (Page, 1986a) and the report for the Wirral peninsula (Page, 1986c). Since no maps were available for the report for Swindon (Page, 1986b), only information detailing the size of the area, boundaries were chosen to encircle the outer urbanised limits of the town, accounting for the urban spread that had occurred in the time between the two surveys.

Within each of the three study areas, visits were made to “high-risk sites” found to provide appropriate conditions for feral cats (Page and Bennett, 1994), including factories and trading estates, academic institutions (schools and colleges), heavy industrial premises, chemical plants, refuse depots, hospitals, docks, derelict land, allotments and plant nurseries. These “high-risk sites” were located based on extensive use of Geographers’ A–Z Street Atlas (Geographers’ A–Z Map Company Limited), local telephone directories, internet searches and GIS mapping. Local knowledge regarding feral cats was also sought from organisations and members of the public who, through the nature of their work, may know the whereabouts of colonies.

Each site was visited and enquiries made to any person(s) who regularly observed the entire site at all times of the day to accurately assess the presence and status of cats on site - for example, security guards. A broad range of questions were asked to determine the number and status of feral cats present, under the definition specified by the Rabies (control) Order 1974: how many cats do you see? How frequently do you see them? How long have they been present on site? What is their condition? Are they neutered or have you seen kittens? Fur patterns were also recorded to reduce the possibility of individuals being counted more than once. Feral cat colonies were then plotted onto maps using ArcView GIS version 3.2 (Environmental Systems Research Institute, 1998).

## 3 Results

### Number of sites surveyed

A total of 741 sites were visited and surveyed for the presence of feral cat colonies in the 1999/2000, compared to 694 in 1986/1987 (Table 1). The changes in the number of “high risk sites” visited reflect the urban development that has occurred in the 12-14 year period between surveys. For example, the number of factories/trading estates and industrial premises has grown by 19.4% since 1987, within the three study areas, leading to 92 more sites of this nature being visited during 1999 and 2000. Dockyards were only found in the Wirral peninsula surveys and, as a result of the recent decline in the ship building industry, two dockyard sites were lost between surveys and were replaced by housing and business development sites respectively.

**Table 1.** The number and location of sites searched for feral cat colonies in the 1986/7 and 1999/2000 surveys of Swindon, Oldham and the Wirral.

| **Type of High Risk Site** | **Swindon** |  | **Oldham** |  | **The Wirral peninsula** |  | **Total** |  |
| --- | --- | --- | --- | --- | --- | --- | --- | --- |
|  | **1986** | **2000** | **1986/7** | **2000** | **1986** | **2000** | **1986/7** | **99/00** |
| Factories / trading estates | 48 | 68 | 295 | 310 | 131 | 188 | **474** | **566** |
| Academic institutions (e.g. schools) | 14 | 10 | 4 | 16 | 30 | 13 | **48** | **39** |
| Heavy Industrial Premises | 2 | 2 | 6 | 3 | 29 | 27 | **37** | **32** |
| Chemical Plant | 0 | 0 | 0 | 1 | 6 | 10 | **6** | **11** |
| Council depots and refuse sites | 12 | 12 | 6 | 4 | 7 | 11 | **25** | **27** |
| Hospitals | 6 | 3 | 3 | 2 | 10 | 6 | **19** | **11** |
| Shops | 0 | 20 | 0 | 0 | 0 | 0 | **0** | **20** |
| Allotments and nurseries | 0 | 1 | 9 | 6 | 14 | 8 | **23** | **15** |
| Hotels | 5 | 2 | 0 | 0 | 0 | 0 | **5** | **2** |
| Docks | 0 | 0 | 0 | 0 | 12 | 10 | **12** | **10** |
| Other | 0 | 0 | 0 | 4 | 0 | 2 | **0** | **6** |
| **Total** | **87** | **118** | **323** | **346** | **239** | **275** | **694** | **741** |

### Number of feral cat colonies identified

The number of feral cat colonies fell from 68 to 43 across all three study areas, representing an overall decrease of 37% between the two surveys (Table 2). The largest decrease was observed in Swindon with 50% fewer colonies, followed by the Wirral peninsula (−43%) and Oldham (−23%). We estimated this decrease in the number of colonies to translate to 212-247 fewer feral cats areas across all three survey sites. Factories/trading estates and industrial premises continue to be the most frequent sites occupied by feral cats. Rises were found in the number of colonies associated with council depots and refuse sites in the Wirral peninsula, and with allotments and nurseries in Oldham, with three colonies located at each site in the recent surveys where previously there were none.

**Table 2.** The number and location of feral cat colonies found in the 1986/7 and 1999/2000 surveys of Swindon, Oldham and the Wirral.

| **Type of High Risk Area** | **Swindon** |  | **Oldham** |  | **The Wirral peninsula** |  | **Total** |  |
| --- | --- | --- | --- | --- | --- | --- | --- | --- |
| **Survey Year** | **1986** | **2000** | **1986/7** | **2000** | **1986** | **2000** | **1986/7** | **99/00** |
| Factories / trading estates | 3 | 3 | 13 | 11 | 17 | 12 | 33 | 26 |
| Institutions | 0 | 0 | 0 | 0 | 0 | 0 | 0 | 0 |
| Heavy Industrial Premises | 0 | 0 | 2 | 0 | 10 | 5 | 12 | 5 |
| Chemical Plants | 0 | 0 | 1 | 0 | 2 | 1 | 3 | 1 |
| Refuse sites | 0 | 0 | 0 | 0 | 0 | 3 | 0 | 3 |
| Hospitals | 3 | 0 | 0 | 0 | 1 | 1 | 4 | 1 |
| Allotments and nurseries | 0 | 0 | 0 | 3 | 0 | 0 | 0 | 3 |
| Docks | 0 | 0 | 0 | 0 | 9 | 0 | 9 | 0 |
| Other | 0 | 0 | 6 | 3 | 1 | 1 | 7 | 4 |
| **Total** | **6** | **3** | **22** | **17** | **40** | **23** | **68** | **43** |

We identified 17 feral cat colonies in Oldham, Greater Manchester, four of which had already been described in the 1987 survey. The colonies show distinct clustering to more developed, urban areas of the town (Figure 2).

**Figure 2:**
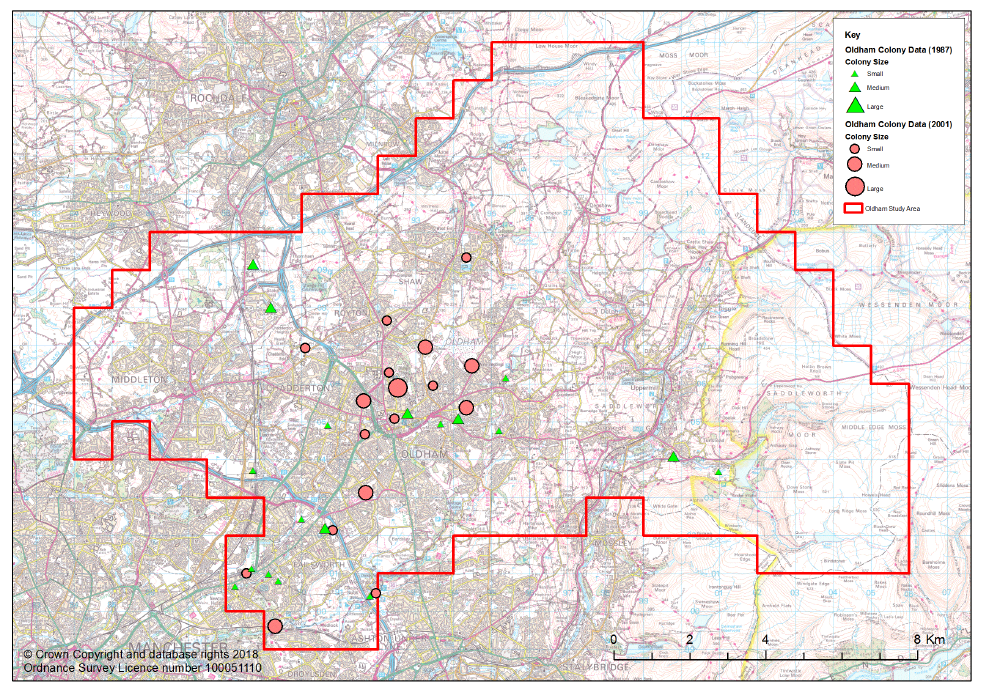
The locations and size of feral cat colonies found in Oldham, Greater Manchester, from surveys carried out in 1987 and 1999-2000. The survey area was 136 km^2^.

The exact locations of the feral cat colonies in Swindon was not recorded in the 1986 survey. However, three colonies described on hospital grounds in 1986 were not present when resurveyed in 2000 (Figure 3).

**Figure 3:**
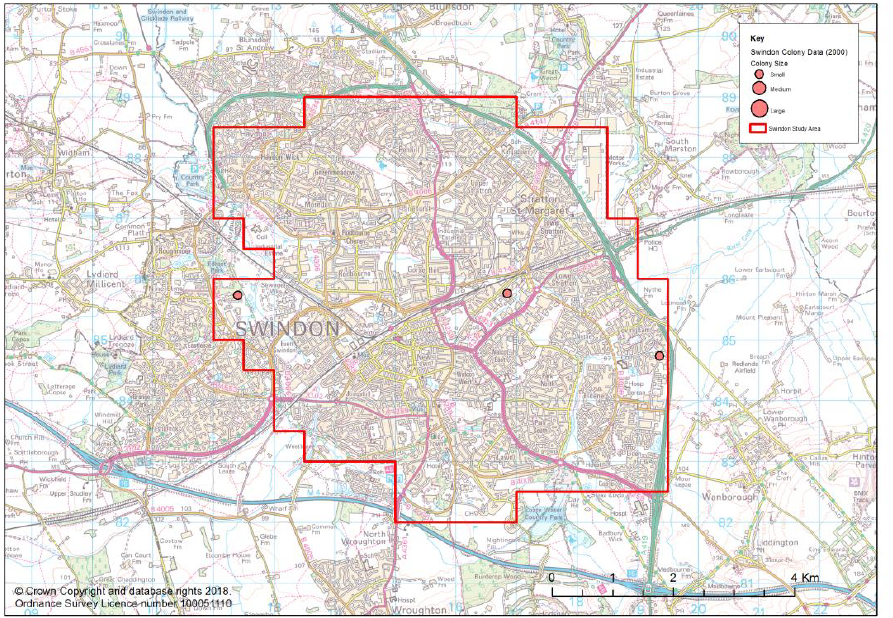
The locations of feral cat colonies from the 2000 survey of Swindon. The survey area was 41.5 km^2^.

We reported 23 feral cat colonies in the Wirral peninsula in 2000, seven of which had already been described in the 1986 survey. A distinct spatial clustering of colonies is noticeable in the more industrial, urban areas along the eastern side of the peninsula (Figure 4).

**Figure 4:**
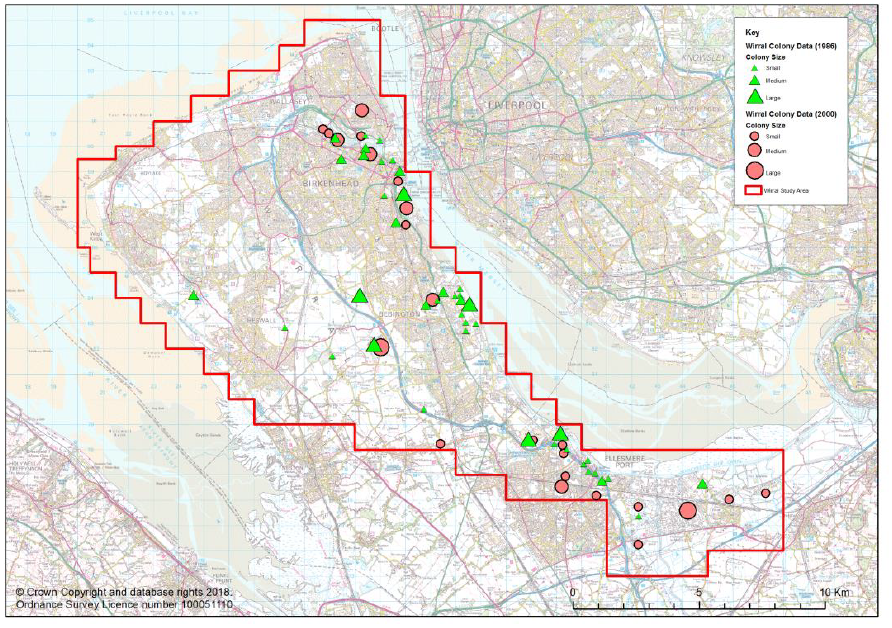
The locations and size of feral cat colonies found in the Wirral peninsula, Merseyside, from surveys completed in 1986 and 2000. The survey area was 228 km^2^.

### Feral cat colony size

The sizes of the colonies identified in both surveys are presented in Table 3.

**Table 3.** The sizes of feral cat colonies (n = number of individuals) found in the 1986/7 and 1999/2000 surveys of Swindon, Oldham and the Wirral peninsula.* no colony sizes were reported for the 1986 Swindon survey.

|  | Year | Small | Medium | Large | Total |
| --- | --- | --- | --- | --- | --- |
| | | $3 \geq n < 5$ | $5 \geq n \leq 15$ | $n > 15$ | |
| Oldham | 1987 | 16 | 6 | 0 | 22 |
|  | 99/2000 | 10 | 6 | 1 | 17 |
| Swindon* | 1986 | ? | ? | ? | 6 |
|  | 2000 | 3 | 0 | 0 | 3 |
| The Wirral | 1986 | 21 | 13 | 6 | 40 |
|  | 2000 | 15 | 7 | 1 | 23 |

Across all three areas, the percentage of medium size colonies (between five and 15 cats) has remained fairly constant between the two surveys (Figure 5). We did however note a halving in the proportion of colonies that were classified as large (>15 cats) with a concomitant small increase in the number of small colonies (<5 cats).

**Figure 5:**
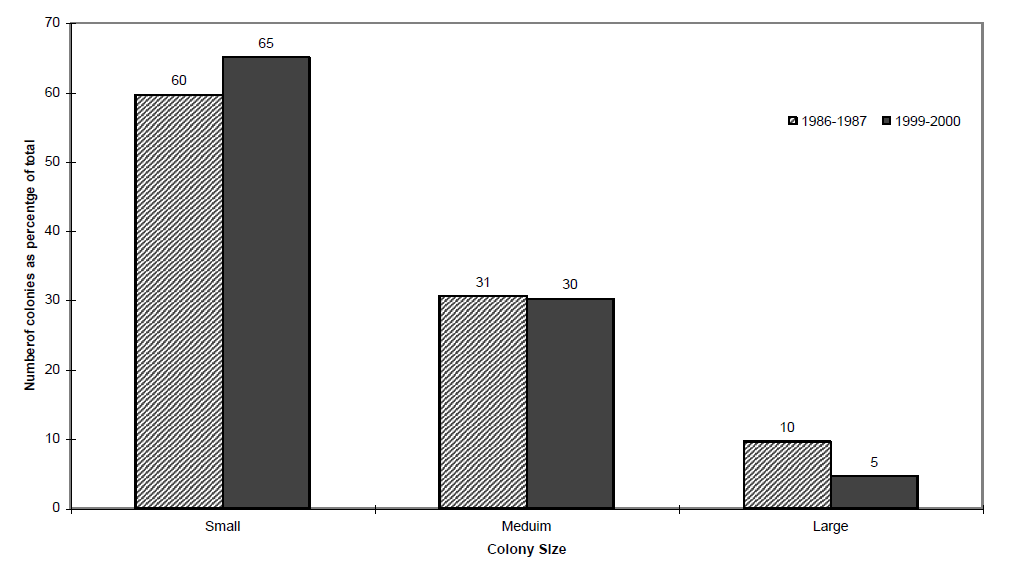
Change in the distribution of the size of feral cat colonies between the 1986/1987 and the 1999/2000 surveys. Small colonies comprise of between three and five cats, medium colonies between five and 15 cats and large colonies harbour >15 cats.

## 4 Discussion

The aim of the study was to re-survey three areas of England to assess the number and distribution of feral cat colonies found there. The results suggest an overall decline in the number of colonies, the number of cats and the size of the colonies between 1986 and 2000. The total number of colonies located during the later survey was 43, compared to 68 previously; a fall of 37%. Whilst enquiring into the presence, status and number of cats at “high risk sites”, information was frequently given for the cause of the reduction in colony size and number of feral cats. The removal of suitable habitat between the two surveys was reported as the main factor contributing to the decline observed in 28% of the sites. This appears to be at odds with our initial observation that the number of potential “high risk sites” had increased from 649 in the 1986/87 survey to 741 in 2000, with a 19% increase in in the number of factories/trading estates and industrial premises, sites found to be most strongly associated with feral cat colonies.

The answer may lie in the type of habitat that has been removed and the recent style of new buildings that have either replaced them, or been built since as new estates. Older buildings tend to be more hospitable to feral cats, providing them with sheltered, quiet places such as ducts, un-used cellars and gaps under buildings, to live and breed in. Heat was provided by boiler rooms or exposed pipes which the cats readily utilised (Dards 1981). Many of these old buildings were demolished, renovated or replaced in more recent years. Modern buildings and industrial estates are more secure and insulated, with minimised loss of heat and energy to the external environment, leaving little or no places for animals to shelter and gain warmth.

Studying feral cats located at Portsmouth docks, Dards (1978, 1981) noted the frequency of usage and availability of steam pipes, providing extensive, covered networks of tunnels used by the cats for shelter and warmth, as well as for moving between areas. These types of habitats were lost as a result of the decline in the ship building industry. Between the two Merseyside surveys, two of the nine docks previously active had been turned into housing or business estates, the remaining seven all suffering from a decline in production.

The declining weaving and textile industry based in Oldham has also had an impact on feral cat habitat. Historically, the mills were places of intense human activity, with feral cats being encouraged to live and breed on site, through the provision of food and shelter by staff, to control rodents. The decline in the number of mills resulted in the loss of 5 colonies, or 38 feral cats, in Oldham. The remaining mills have for the most part been repurposed into food production sites (on which all animals had to be legally removed by pest control companies) or business centres which do provide some suitable conditions for feral cats; three colonies being found in 1999/2000 at such premises.

Many of the factories/trading estates and industrial premises visited carried out some form of pest control at frequent intervals. Although control is infrequently targeted directly at feral cats, secure bait containers are often placed down containing poison for rodents, which reduces the amount of food for feral cats.

The present survey confirmed that the concept of confining visits during a rabies contingency to “high risk sites” is an efficient method for finding any feral cat colonies (Rees, 1981; Page and Bennett, 1994). Of the colonies located in the more recent survey 91% were found to be associated with these sites. Factories, trading and industrial estates remain the most common type of site found to be occupied by feral cats. Three colonies were found in 1999/2000 at council depots/refuse sites, and allotments/ plant nurseries where previously none were associated with such areas. Despite 87 academic institution sites being visited across both surveys, no feral cat colonies were recorded suggesting that this type of site is not offers unsuitable conditions for feral cats and that they should be removed from the list of “high risk sites” thus reducing the time and effort required to survey an area during a rabies outbreak. The four colonies found outside of “high risk sites” were located through the knowledge of local animal welfare groups. We encountered difficulties entering survey sites on only on two occasions, out of 741 premises visited. These were sites with high levels of security, a military base and large oil refinery. At present, the mean density of feral cats colony is one colony per 9.43 km^2^, falling from a mean of one colony per 5.96 km^2^ in 1987. Combining the results of all the studies past and present suggest on average one person can survey an area of 6.1 km^2^ per day (n = 6, standard deviation = 3.7).

The definition of a feral cat colony in the context of rabies contingency purposes, i.e. cats which the owner of the land has no legal obligation to confine, proved to be somewhat difficult to apply in the field. Any cats falling out of this definition during a rabies control operation should be confined by the person(s) responsible for them. Many cat colonies were wanted for one reason or another, whether to control rodents or for the pleasure of cat lovers on a particular site, however by law, these people would not be obliged to confine such animals. Some colonies were found to be located over several sites with as many feeders. Cats found to be free-roaming and unconfined at all times (with no more responsibility taken by the owner/occupier than feeding) were considered feral during this survey, which through inquiries was considered the same in the previous study by MAFF staff. Groups of cats found around single houses in residential areas, where the cats lead a very domesticated life-style, were considered non-feral as a result of a person’s high level of responsibility and interaction with the animals.

We recommend that feral cat surveys should be repeated to determine up to date feral cat population and density data.

## 5 Acknowledgments

We are grateful to all the people approached during the study, and to David Fouracre for production of the figures.

